# Anatomo-functional changes in neural substrates of cognitive memory in developmental amnesia: Insights from automated and manual MRI examinations

**DOI:** 10.1101/2023.01.23.525152

**Authors:** Loïc J. Chareyron, W.K. Kling Chong, Tina Banks, Neil Burgess, Richard C. Saunders, Faraneh Vargha-Khadem

## Abstract

Despite bilateral hippocampal damage dating to perinatal or early-childhood period, and severely-impaired episodic memory that unfolds in later childhood, patients with developmental amnesia continue to exhibit well-developed semantic memory across the developmental trajectory. Detailed information on the extent and focality of brain damage in these patients is needed to hypothesize about the neural substrate that supports their remarkable capacity for encoding and retrieval of semantic memory. In particular, we need to assess whether the residual hippocampal tissue is involved in this preservation, or whether the surrounding cortical areas reorganise to rescue aspects of these critical cognitive memory processes after early injury.

We used voxel-based morphometry (VBM) analysis, automatic (FreeSurfer) and manual segmentation to characterize structural changes in the brain of an exceptionally large cohort of 23 patients with developmental amnesia in comparison with 32 control subjects.

Both the VBM and the FreeSurfer analyses revealed severe structural alterations in the hippocampus and thalamus of patients with developmental amnesia. Milder damage was found in the amygdala, caudate and parahippocampal gyrus. Manual segmentation demonstrated differences in the degree of atrophy of the hippocampal subregions in patients. The level of atrophy in CA-DG subregions and subicular complex was more than 40% while the atrophy of the uncus was moderate (−23%). Anatomo-functional correlations were observed between the volumes of residual hippocampal subregions in patients and selective aspects of their cognitive performance viz, intelligence, working memory, and verbal and visuospatial recall.

Our findings suggest that in patients with developmental amnesia, cognitive processing is compromised as a function of the extent of atrophy in hippocampal subregions, such that the greater the damage, the more likely it is that surrounding cortical areas will be recruited to rescue the putative functions of the damaged subregions. Our findings document for the first time not only the extent, but also the limits of circuit reorganization occurring in the young brain after early bilateral hippocampal damage.

## Introduction

In humans, hypoxic-ischaemic events experienced early in life can damage specific subcortical brain structures such as the hippocampus, the basal ganglia, and the thalamus^1–6^. White matter abnormalities have also been found in newborns with hypoxic-ischaemic encephalopathy^7–9^, and these abnormalities can persist into adolescence^10^.

We previously identified a group of patients with hippocampal atrophy who had suffered hypoxic-ischaemic episodes in infancy or childhood and later developed a memory disorder differing from that commonly described in patients with adult-onset amnesia of temporal-lobe origin^4,11^. In the latter cases, the disorder most often takes the form of a global anterograde memory loss (i.e. one affecting both episodic and new post-injury semantic memory, thus severely restricting both recall and recognition processes)^12–15^. In patients with hippocampal damage of early-onset, by contrast, the disorder is more limited, being characterized by markedly impaired episodic memory, recall/recollection, spatial processing and navigation deficits, but relatively preserved semantic memory, working memory, and recognition performance. We have labelled this dissociated form of memory function ‘developmental amnesia’^6,11,16–18^. Given the early onset of hippocampal damage, the compensatory reorganization that has occurred in patients with developmental amnesia is likely to be distinct from what is observed in patients with hippocampal damage acquired in adulthood. The differential maturation of distinct hippocampal circuits during normal development might underlie the differential emergence of specific hippocampus-dependent memory processes^19^. In patients with adult-acquired hippocampal lesions, the damage has occurred to already-established, and normally-functioning memory circuits, whereas in patients with developmental amnesia, the early bilateral lesions have probably led to the development of a differently-organized memory system. For these reasons, it is difficult, and somewhat inappropriate, to compare memory function and cognitive profiles in patients with developmental amnesia to those of patients with adult-onset amnesia^20^.

Among the many questions that have arisen from the study of patients with developmental amnesia, the foremost concerns the integrity of other regions of the brain, particularly the medial temporal lobe (MTL) and/or preserved hippocampal subregions that could be involved in memory circuit reorganization following early hippocampal damage.

The degree of hippocampal atrophy in patients with developmental amnesia ranges from 28% to 62% compared to healthy controls^4^. This large range of hippocampal volume reduction in this cohort suggests that there may indeed be variability in the hippocampal response to hypoxia-ischemia at the level of hippocampal subregions. Hypoxia-ischemia is known to differentially affect the hippocampal fields and subdivisions. Studies of animal models of cerebral ischemia have reported that a brief episode results in selective neuronal death in the CA1 field of the hippocampus^21,22^ while the adjacent CA3 remains less vulnerable^23^. In humans, while damage to CA1 is consistently reported following cerebral ischemia, damage to additional hippocampal regions is much more variable^24^.

Postmortem examinations have reported some neural cell loss in layers of entorhinal cortex in patients who suffered ischemic episodes in adulthood^25^. However, it was reported that the volume of the parahippocampal gyrus was not significantly reduced in patients with adult-onset amnesia presenting with bilateral hippocampal damage^26^. A cortical thickness analysis performed in a single patient with developmental amnesia didn’t find abnormality in the subhippocampal structures within the medial temporal lobe, including the perirhinal and ventral entorhinal cortices^27^. Also, a recent study demonstrated that parahippocampal activity during scene reinstatement in patients with developmental amnesia was similar to controls^28^. However, the structural integrity of the perihippocampal gyrus, and by implication, the integrity of the hypothesized substrate of semantic memory and other preserved mnemonic processes remain to be quantitatively assessed in patients with developmental amnesia.

Here, we studied the largest cohort of patients with the rare condition of developmental amnesia ever assembled to document the extent of structural brain damage and hypothesize possible circuit reorganizations that may be involved in compensating hippocampal function. We used voxel-based morphometry (VBM) and FreeSurfer automated segmentation on MRI scans to estimate the extent of tissue changes in brain regions. We also used manual segmentation to estimate the volume of three hippocampal subregions (uncus, CA-DG and subicular complex) and three surrounding cortical areas (entorhinal, perirhinal and parahippocampal cortices) in patients with developmental amnesia and healthy controls. Controls and patients were assessed with neuropsychological tests of intelligence (WISC or WAIS), and recall/recognition for verbal and visual material (Doors and People Test). Finally, we investigated the relationships between degree of atrophy of hippocampal subregions and cognitive deficits.

## Materials and methods

### Participants

The patients with developmental amnesia were characterized by (i) a history of episodes of hypoxia-ischemia in early life, (ii) quantified hippocampal volume reduction above 25% relative to a group of controls, and (iii) severely impaired episodic memory^4^. The most common aetiology was perinatal hypoxic-ischemic encephalopathy (Supplementary Table 1). While long-term neurodevelopmental outcome depends on the severity of the hypoxic-ischemic episode, subtle cognitive deficits and behavioural alterations have been revealed through long-term evaluations even in mild forms of hypoxia-ischemia^29^.

When all the manual volume estimates were completed, one of the patients, who presented with moderate memory deficits and only 11.0% hippocampal volume reduction associated with two unconfirmed episodes of prolonged seizures at age 4, was excluded from the developmental amnesia group.

Also, we excluded from the developmental amnesia group a patient with temporal lobe epilepsy presenting with a 25.8% hippocampal volume reduction, but without a documented episode of early hypoxia-ischemia and/or status epilepticus. Additionally, blinded neuroradiological examination carried out independently by one of the authors (WKC) revealed that this patient’s MRI scan was entirely normal. The hippocampus, fornix and mammillary bodies were noted to have a normal appearance in contrast to all other patients with developmental amnesia.

The developmental amnesia group included three patients (DA05, DA06, DA10) who had sustained hypoxic–ischemic episodes between the ages of 9 and 12 years in contrast to other patients who had suffered such episodes perinatally or within the first 3 months of life. However, it has been previously demonstrated that the early (below age one), and late (between 6-14 years) groups exhibit similar profiles, and thus concluded that the effective age at injury for developmental amnesia extends from birth to puberty^30^.

Voxel-based morphometry (VBM), automatic (FreeSurfer) and manual segmentations were performed on 32 control participants (16 females; mean age: 17.7 years, SD 7.7, range 8−38) and 23 patients with developmental amnesia (11 females; mean age: 19.5 years, SD 9.0, range 8−40).

The study was approved by the London-Bentham Research Ethics Committee (REC Reference No. 05/Q0502/88) and all participants and their carers read an age-appropriate information sheet and gave written informed consent according to the Declaration of Helsinki, before commencing the study.

### Structural MRI

Participants were either scanned with a 1.5T-MRI scanner (24 control participants, 16 patients with developmental amnesia) or with a 3T-MRI scanner (8 control participants, 7 patients with developmental amnesia).

Whole brain structural 1.5T-MRI scans were obtained using a Siemens Avanto Scanner, with a T1-weighted 3D FLASH sequence with the following parameters: repetition time 11 ms, echo time 4.94ms; in-plane resolution 1 mm × 1 mm; slice thickness of 1 mm^31^.

Whole brain structural 3T-MRI scans were obtained using a 3T Siemens MRI system with a 20 channel head coil. A T1-weighted magnetization-prepared rapid gradient-echo (MPRAGE) scan was acquired with the following parameters: in-plane resolution of 1 mm × 1 mm; slice thickness of 1 mm; repetition time of 2,300 ms; echo time of 2.74 ms^32^.

### VBM analysis

Processing of the structural MRI scans was carried out using Statistical Parametric Mapping 12 (SPM12^33^; version 7771, downloaded from https://www.fil.ion.ucl.ac.uk/spm/) running on MATLAB 2020b (The Mathworks, Natick, MA, USA). Using the segmentation procedure implemented in the CAT12 toolbox (https://neuro-jena.github.io/cat//index.html), the images were segmented into grey matter (GM), white matter (WM) and cerebrospinal fluid (CSF). For each participant, this resulted in a set of 3 images in the same space as the original T1-weighted image, in which each voxel was assigned a probability of it being GM, WM and CSF. The CAT12 toolbox was also used to ensure the quality of the segmentation, which was either ‘good’ (‘B’, controls *n* = 2; patients *n* = 1), ‘satisfactory’ (‘C’, controls *n* = 25; patients *n* = 17) or ‘sufficient’ (‘D’, controls *n* = 5; patients *n* = 5). The structural MRI data were analysed using the smoothed, modulated grey matter segments from 23 patients and 32 controls. A general linear model with factors group (control, patients) and covariate of no interest (intracranial volume) was used to identify the group-wise pattern of GM atrophy in patients with developmental amnesia. Statistical maps of volume change were displayed using a voxel-wise threshold of *P* < .001. The Anatomy toolbox for CAT12 was used to determine the probability of the voxels being located in particular structures (e.g. hippocampus).

### FreeSurfer analysis

Cortical reconstruction and volumetric segmentation was performed with the FreeSurfer software v7.3.2 (downloaded from https://surfer.nmr.mgh.harvard.edu/). Briefly, this processing includes motion correction and averaging of multiple volumetric T1 weighted images, removal of non-brain tissue, segmentation of the subcortical white matter and deep gray matter volumetric structures, intensity normalization, and parcellation of the cerebral cortex into units with respect to gyral and sulcal structure producing representations of cortical thickness^34–36^. We also used a FreeSurfer tool that produces an automated segmentation of the hippocampal substructures^37^.

### Manual segmentation

For the manual measurement of regions of interest (ROI) volumes, the data were reformatted into 1 mm-thick contiguous slices in the coronal plane. ROI cross-sectional areas were measured in all slices along the entire length of the hippocampus, entorhinal, perirhinal and parahippocampal cortices using ITK-SNAP (version 3.8.0; www.itksnap.org)^38^. Manual tracing is still the gold standard for measuring hippocampal volume and is particularly appropriate for samples of relatively small size,^39^ presenting with various levels of hippocampal atrophy,^40^ or including children and adolescents^41^. All manual measurements of the ROIs were carried out by one of the authors (LJC) who remained blind to participant identity (Supplementary Fig. 1-2).

The hippocampus comprises the CA fields (CA3, CA2, CA1), the dentate gyrus (DG), subiculum, presubiculum, and parasubiculum^42^. The human uncus [also termed the “uncal portion of the hippocampal formation”]^43^ is situated antero-medially and consists of slightly modified DG, CA3, CA2, CA1, and subiculum^44^. For the segmentation of the hippocampus and its subregions, we followed the guide provided by Dalton *et al*.^45^. Given the difficulty to distinguish hippocampal CA fields on 1.5T and 3T-MRI acquisitions, we restricted our volumetric analyses to three ROIs: uncus [defined according to Ding and Van Hoesen]^44^, CA-DG (CA fields and dentate gyrus), and subicular complex (subiculum, presubiculum, and parasubiculum). The segmentation of the entorhinal, perirhinal and parahippocampal cortices was based on structural descriptions of human medial temporal lobe and segmentations protocols^46–50^.

The volumes in cubic millimetres were calculated by summing the number of voxels (1×1×1 mm) for each ROI. The volume of the hippocampus as a whole was calculated as the sum of hippocampal subregions’ volumes (uncus, CA-DG, subicular complex). We then used parametric tests as volumetric estimates of the hippocampal subregions and cortical areas were normally distributed (Shapiro-Wilk normality test; control: all *P* > .06; patients: all *P* > .09). The volume of all the ROIs was calculated as the mean of the left and right hemisphere measurements.

Intracranial volume (ICV) was manually measured on the sagittal dataset with a 1-in-10 random and systematic sampling strategy. The ICV was normally distributed (Shapiro-Wilk normality test; control: *P* = .05; patients: *P* = .23). The ICV was 6.8% smaller in the developmental amnesia group than in the control group (t_(53)_ = −2.70; *P* = .009). Corrections, derived from the regression line of control ROI volume (V) versus ICV, were then made for ICV based on the formula [ΔV = a × (ΔICV)] where ‘a’ is the regression coefficient.

### Behavioural analyses

#### WISC/WAIS

The Wechsler Adult Intelligence Scale, 3rd Ed. (WAIS-III) and the Wechsler Intelligence Scale for Children, 4th Ed. (WISC-IV) provide four index scores: Verbal Comprehension Index, measuring understanding and expression of verbal concepts; Perceptual Reasoning Index, reflecting visuospatial perception and perceptuomotor manipulation; Working Memory Index, tracking on-line immediate memory and executive control; and Processing Speed Index, measuring speed of visuoperceptual discrimination, and copying of number-symbol associations. The four index scores are expressed as standard scores with a mean of 100 and a standard deviation of 15.

##### Scoring

The WISC-IV or WAIS-III were administered to all patients with developmental amnesia (*N* = 23, 11 females, mean age: 19.5 years, SD: 9.0, range: 8−40) and to 23 controls (13 females, mean age: 18.0 years, SD: 8.4, range: 8.2–38). For the analysis of the data, we then used parametric tests as the data were normally distributed in all four indexes (Shapiro–Wilk test of normality; Control: all *P* > .36; patients: all *P* > .10).

#### The Doors and People Test

The Doors and People Test was administered according to the instructions in the published manual,^51^ as described in detail in previous reports^16,52^. Briefly, for the verbal recall subtest (‘People’), participants were presented with four pictures of people and after three learning trials asked to recall their names cued by their profession. For the visual recall subtest (‘Shapes’), participants first copied, and drew the shapes from memory after three learning trials. For verbal recognition (‘Names’), participants were presented with a list of names, and then asked to recognize each one from a list of four alternatives. Finally, in the visual recognition subtest (‘Doors’), participants viewed a list of doors and then identified each one from a list of four alternatives. See Supplementary Information for more details.

##### Scoring

The Doors and People Test was administered to all patients with developmental amnesia (*N* = 23, 11 females, mean age: 19.5 years, SD: 9.0, range: 8−40) and to 21 controls (11 females, mean age: 17.6 years, SD: 8.6, range: 8−38). One of the patients performed all but the ‘Names’ subtest (‘People’, ‘Doors’, ‘Shapes’: *N* = 23; ‘Names’: *n* = 22). Given that the developmental amnesia group included individuals under the age of 16, which is the first age band at which standard scores are available on the Doors and People Test, we used standard scores based on the scores of our control group; raw scores (SD): People, 28.2 (6.9); Doors, 18.9 (3.8); Shapes, 34.6 (2.7); Names, 18.6 (3.7). We converted patients’ and controls’ raw scores to z-scores relative to the control group’s scores. For the analysis of the data, we then used nonparametric tests as the data were not normally distributed in all four subtests (Shapiro–Wilk test of normality; Control: *P* < .001; patients: *P* = .044).

## Results

### Voxel-based morphometry

CAT12/SPM12 voxel-based morphometry analysis provided estimates of grey matter (GM) volumes in patients with developmental amnesia and controls (Fig 1,2). GM volume was significantly reduced bilaterally in the hippocampus (–29.8% ; *P* < .0001 after Bonferroni correction), thalamus (–22.3% ; *P* < .0001), and amygdala (–17.5% ; *P* < .0001) in patients with developmental amnesia compared to controls. Milder GM volume reductions were observed in patients with developmental amnesia in the left caudate nucleus (–14.7% ; *P* = .013) and in the left parahippocampal gyrus (–12,6% ; *P* = .047). CAT12/SPM12 also provided estimates of cortical surfaces (thickness) based on different atlas maps. Analysis based on the ’Destrieux’ cortical atlas^53^ didn’t reveal any significant thickness reduction in the cortex of patients with developmental amnesia compared to controls (after Bonferroni correction). However, analysis based on the gyral-based atlas ’Desikan-Killiany-Tourville’^54^ revealed a mild 9.3% reduction in the left parahippocampal gyrus thickness in patients with developmental amnesia (P = .015).

**Figure 1.**
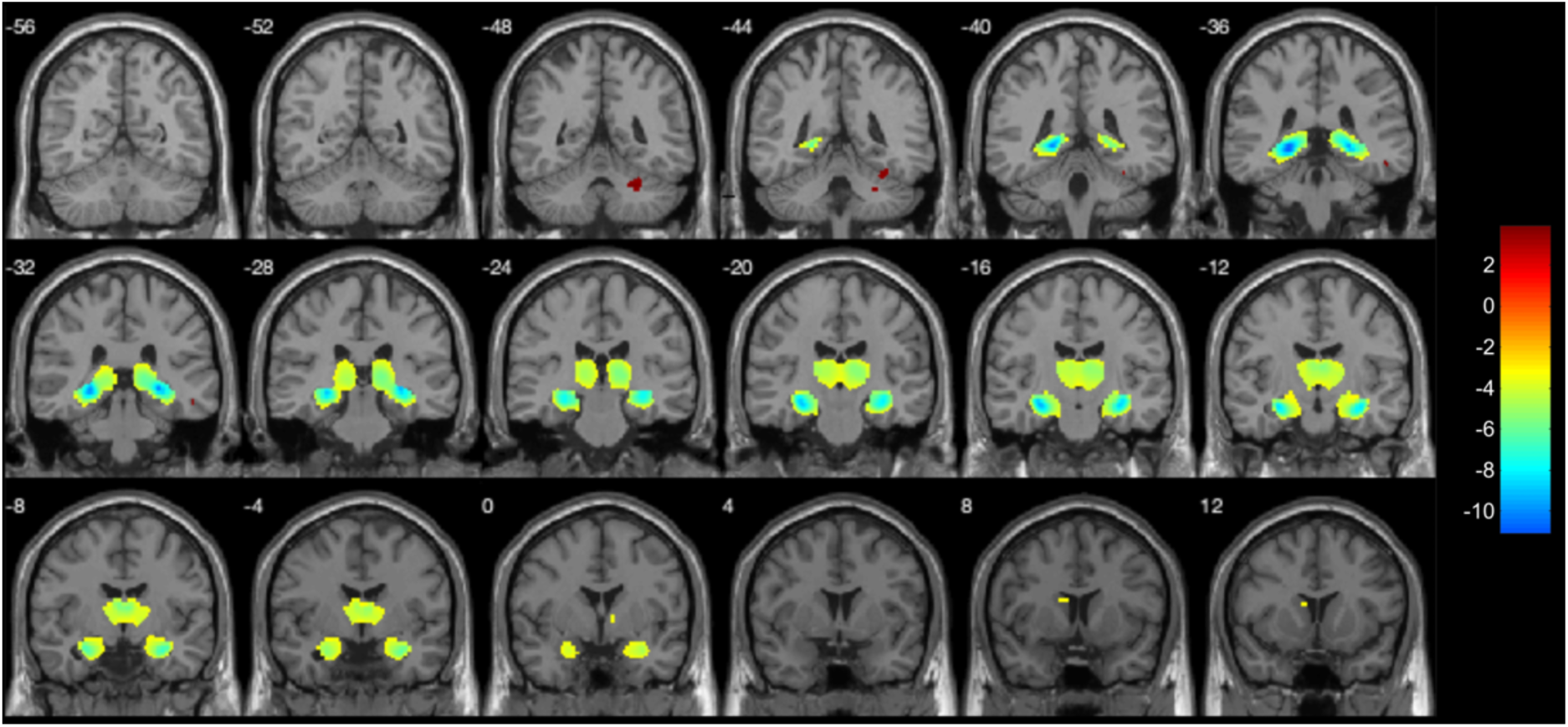
Results of CAT12/SPM12 voxel-based morphometry analysis comparing patients with developmental amnesia to healthy controls. Differences between a group of controls (*N* = 32) and a group of patients with developmental amnesia (*N* = 23). Main differences were observed in the bilateral hippocampus, thalamus and amygdala (*P* < .001).

**Figure 2.**
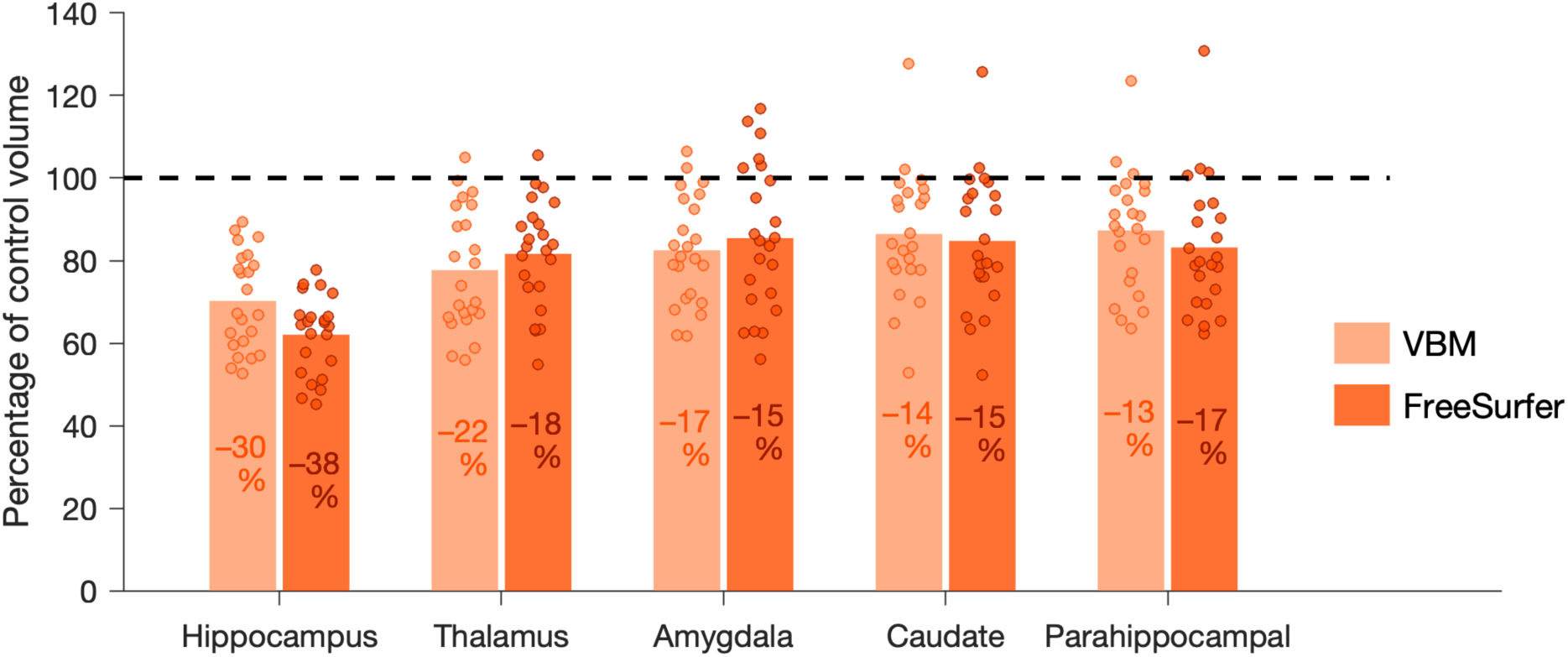
Brain areas with significant volume reduction in patients with developmental amnesia according to both voxel-based morphometry (VBM) and FreeSurfer analysis. Volumes of ROI in patients with developmental amnesia (*N* = 23) are shown as a percentage of the mean control volume (*N* = 32). Percentage values refer to the mean difference to control group. Volumes are calculated as the average of left and right hemisphere volumes.

### FreeSurfer automatic segmentation

FreeSurfer provided volume estimates of subcortical areas in patients with developmental amnesia and controls (Fig 2). Significant volume reductions were observed bilaterally in patients with developmental amnesia in the hippocampus (left: –36.4% ; right: –39.4% ; both *P* < .001 after Bonferroni correction), thalamus (left: –18.6% ; right: –18.1% ; both *P* < .001), amygdala (left: –13.6% ; right: –15.5% ; both *P* < .05), caudate (left: –16.2% ; right: –14.3% ; both *P* < .05) and putamen (left: –13.9% ; right: –11.9% ; both *P* < .05). FreeSurfer identified reduction in the cerebral white matter volume in patients with developmental amnesia (left: – 12.0% ; right: –10.6% ; both *P* < .05). FreeSurfer also provided estimates of cortical GM volumes. Significant cortical GM volume reduction was observed in patients with developmental amnesia in the right parahippocampal cortex only (–17.8% ; *P* < .001 after Bonferroni correction).

Automated segmentation of hippocampal substructures using the FreeSurfer tool revealed more pronounced atrophy of the posterior hippocampus (body + tail) than the hippocampal head in patients with developmental amnesia (–40% and –33% respectively ; paired t test, t_(22)_ = –3.81 ; *P* < .001).

### Manual segmentation estimates

Three hippocampal subregions (uncus, CA-DG and subicular complex) and three surrounding cortical areas (entorhinal, perirhinal and parahippocampal cortices) were manually segmented in healthy controls (*N* = 32) and developmental amnesia patients (*N* = 23) (Supplementary Fig. 1-2). The volume of the hippocampus as a whole was 40% smaller in the patients than in the control group (range: −26% to −53%; t_(53)_ = −15.32; *P* < .001) (Fig. 3; Supplementary Table 2). The volume of the hippocampal subregions was also significantly smaller in the patients than in the control group (2-way ANOVA, F_(1,159)_ = 392.5, *P* < .001). The uncus was 23% smaller (range: +4% to −49%; Tukey’s HSD post-hoc test: *P* = .037), the CA-DG subregions were 45% smaller (range: −28% to −59%; *P* < .001), and the subicular complex was 41% smaller (range: −22% to −56%; *P* < .001). The level of atrophy of the patients’ hippocampal subregions differed significantly from each other (F_(2,66)_ = 23.0, *P* < .001). Tukey’s HSD post-hoc test indicated that the CA-DG and subicular complex showed greater atrophy than the uncus (both *P* < .001). Atrophy of the hippocampus in the patients group was significantly lower in its anterior (−36%) than its posterior (−44%) segment (using the uncal apex as a landmark ; paired t test, t_(22)_ = 2.72, P = .012). Because no differences were observed between the anterior and the posterior segment of the CA-DG (paired t test, t_(22)_ = 0.94, P = .36) or subicular complex (t_(22)_ = 1, P = .33) subregions, the lower level of atrophy observed in the anterior part of the whole hippocampus can be attributed to the relatively more preserved uncus.

**Figure 3.**
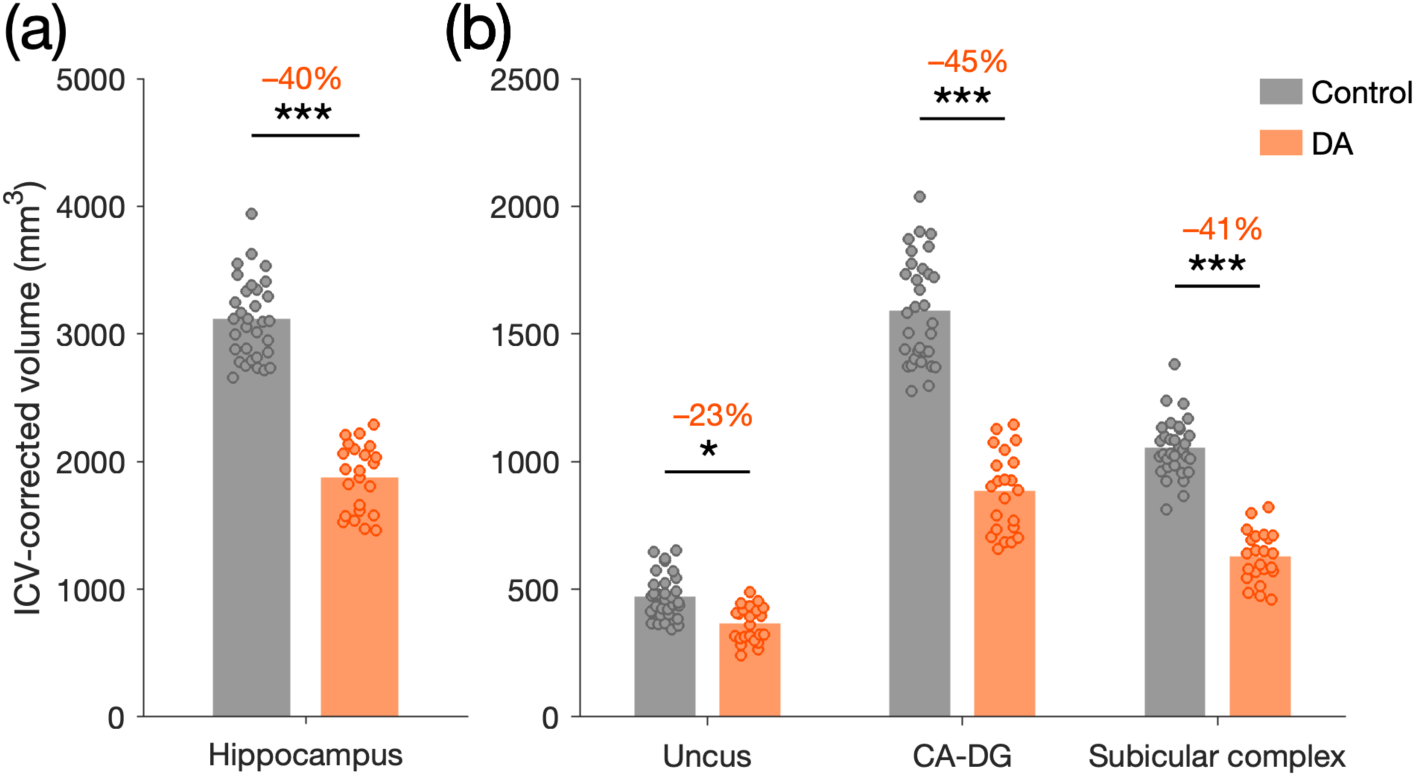
Volume of the hippocampus and its subregions in developmental amnesia. Volume of the hippocampus (**a**) and its subregions (**b**, uncus, CA-DG, and subicular complex) in control (grey; *N* = 32) and developmental amnesia (orange; *N* = 23) groups. Data are represented as mean ± SD. Percentage values refer to the mean difference to control group. Volumes are calculated as the average of left and right hemisphere volumes. All volumes are corrected for intracranial (ICV) volume. *: *P* < .05; ***: *P* < .001.

There was a global difference in the volume of the entorhinal, perirhinal and parahippocampal cortices between the control and patient groups (Supplementary Table 2; 2- way ANOVA, F_(1,159)_ = 6.7, *P* = .010), but no statistical differences in pairwise comparisons (Tukey’s HSD, all *P* > .35).

### Comparison between segmentation methods

Hippocampal volume estimates obtained with the VBM, FreeSurfer, and manual segmentation protocols were compared. While the automated methods reported reliable group differences, VBM and FreeSurfer overestimated the hippocampal volume by 36% and 12% respectively compared to manual segmentation. The correlations between the hippocampal volume estimates obtained by the different methods were poor, especially in patients with developmental amnesia (Supplementary Fig. 3).

### Neuroradiological assessment

Neuroradiological ratings based on visual inspection of the MR images provided independently by one of the authors (WKC) revealed abnormally small fornix and mammillary bodies in all but two patients with developmental amnesia (Supplementary Table 1). No abnormalities were detected in the parahippocampal gyrus (perirhinal, entorhinal, and parahippocampal cortices), or the thalamus, and the basal ganglia. Additional visible abnormality was observed in some cases in the white matter and the cerebellum. The pattern of visible abnormality detected within the hippocampal circuit, and the notable absence thereof in other cortical and subcortical structures, was consistent across individual patients irrespective of their varying aetiology, and age at hypoxic-ischaemic-induced hippocampal damage.

### Doors and People Test

All patients with developmental amnesia (*N* = 23) and 21 of the control participants were assessed on the Doors and People Test, which provides equated measures of recognition and recall in the visual and verbal domains. All patients’ subtest scores were significantly below zero (Wilcoxon signed rank test: all Z < −2.4, all *P* < .016), indicating that patients were impaired relative to controls (who represent zero, as their scores were used to convert raw scores into z-scores). The patients’ scores for the four Doors and People subtests differed significantly from each other (Friedman’s test: χ^2^_(3,n=22)_ = 43.7, *P* < .001). The visual recall (‘Shapes’) subtest yielded a significantly greater deficit than all of the other subtests (all *P* < .001). The degree of deficit on the verbal recall (‘People’) subtest was greater than on the two recognition subtests (both *P* < .02). The degree of deficit was greater on the visual recognition (‘Doors’) subtest than on the verbal recognition (‘Names’) subtest (P = .001). Collapsing across subtests, we found a greater deficit in recall compared with recognition (Z = −4.04; *P* < .001), and a greater deficit in memory for visual compared with memory for verbal material (Z = −4.11; *P* < .001).

We observed significant negative correlations between the two recall subtest scores and the volume of the uncus in the developmental amnesia group [Verbal recall (‘People’): Pearson’s r_(22)_ = –0.47; *P* = .02; Visual recall (‘Shapes’): r_(22)_ = –0.55; *P* = .007; Fig. 4ab]. No other correlations for other subtests or other regions of interest in the developmental amnesia group were significant (all *P* > .09; Supplementary Table 3). Collapsing across test material confirmed that scores on recall were inversely correlated with the volume of the uncus in the developmental amnesia group (r_(22)_ = –0.58; *P* = .004; Fig. 4c). As a corollary, we observed that patients with more preserved uncus had significantly lower scores on recall than patients with more damaged uncus (Z = −2.74; *P* = .006). In the control group, no correlations for any subtests or regions of interest were significant (all *P* > .11; Supplementary Table 3).

**Figure 4.**
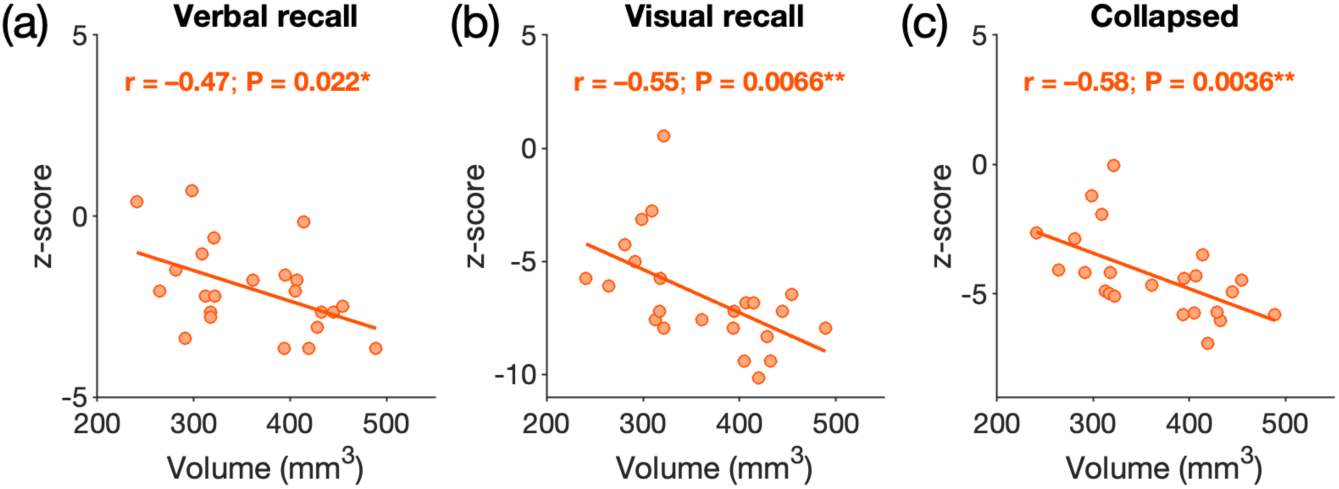
Doors and People recall scores correlations with the volumes of the uncus in patients with developmental amnesia. The patients’ scores (N = 23) on recall subtests of the Doors and People Test correlated negatively with the volume of the uncus. (**a**) Verbal recall (‘People’) subtest. (**b**) Visual recall (‘Shapes’) subtest. (**c**) Collapsed scores. Pearson’s correlation coefficient. All volumes are corrected for intracranial (ICV) volume. *: *P* < .05; **: *P* < .01.

### WISC/WAIS

All patients with developmental amnesia (*N* = 23) and 23 of the control participants were assessed on the WISC or WAIS. Patients were impaired relative to our group of controls only for the Perceptual Reasoning Index scores (2-way ANOVA, F_(1,176)_ = 10.61, *P* = .001; Tukey’s HSD post-hoc test, *P* = 0.04; all other *P* > .25). There were significant inverse correlations between the volume of the uncus and patients’ scores for the Verbal Comprehension Index (VCI), the Working Memory Index (WMI) and the Processing Speed Index (PSI) (all r_(22)_ < – 0.42 ; all *P* < .05; Fig. 5; Supplementary Table 4). No other correlations for other ROIs in the developmental amnesia group were significant. In controls, the WMI scores were positively associated with the volume of the subicular complex (r_(22)_ = 0.46 ; *P* = .029).

**Figure 5.**
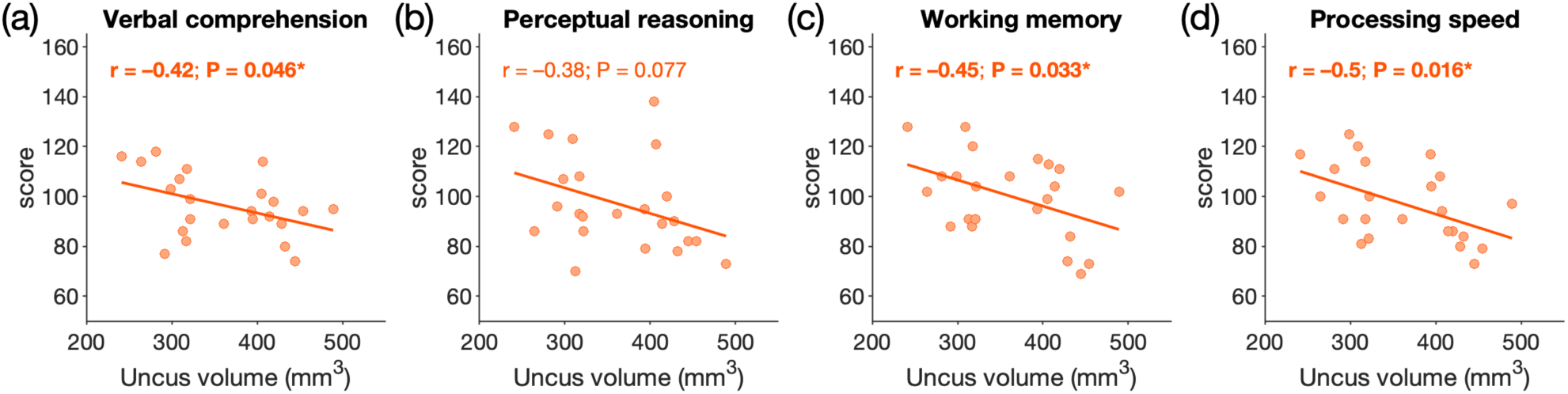
Correlations between WISC/WAIS indexes scores and the volume of the uncus in patients with developmental amnesia. Patients’ scores on Wechsler Intelligence Scales WISC or WAIS (N = 23). (**a**) Verbal comprehension Index; (**b**) Perceptual Reasoning Index;. (**c**) Working Memory Index; (**d**) Processing Speed Index. Pearson’s correlation coefficient. All volumes are corrected for intracranial (ICV) volume. *: *P* < .05.

## Discussion

In this study, we documented the extent of brain damage in patients who had suffered hypoxic-ischaemic episodes in infancy or childhood and later developed the syndrome of developmental amnesia. While the hippocampus exhibited the greatest degree of atrophy, we show that damage also occurs in the thalamus and amygdala, and to a lesser extent in the caudate and parahippocampal gyrus in patients with developmental amnesia. These findings replicate and extend previous reports in the same cohort of patients ^4,5^. We also addressed the question of whether the volumes of residual hippocampal subregions are associated with cognitive memory performance in these patients. Using a combination of neuropsychological test results, we showed several inverse relationships between the volume of the residual uncus and patients’ performance on recall memory, Verbal comprehension Index, Working memory Index and Processing speed Index. The effects reported here suggest that in patients with developmental amnesia, the remaining hippocampal tissue, depending on its extent of damage, provides signals that may inhibit compensatory mnemonic development by the relatively preserved surrounding cortical areas. This process could occur either because normally silent connections promote compensation, or because signals (either typical or pathological) inhibit compensation. However, whatever the extent of hippocampal damage in developmental amnesia, and despite these compensatory reorganizations, hippocampal-dependent episodic memory retrieval remains irreversibly compromised.

### Methodological considerations

Although manual tracing is still the gold standard for measuring the volumes of hippocampal subregions and cortical areas on MRI acquisitions, especially in the case of patients, manual segmentation is subjective and susceptible to measurement errors. Also, the precision of the measure is limited by the resolution of the MRI scans. However, analyses of MR images for estimating the extent of hippocampal damage in amnesic patients have been validated by postmortem examinations^25^. In particular, a clear concordance has been reported in the appearance of the damaged hippocampus on the MRI and in the histological sections^25^.

Here, we used in addition a combination of automated and manual approaches, all of which showed the same pattern of damage in patients, particularly along the antero-posterior axis of the hippocampus and the parahippocampal gyrus. However, the present study shows that while automated methods can reliably detect differences between experimental groups, hippocampal volume estimates obtained with automated methods are overestimated and cannot be correlated with manually obtained volume estimates, highlighting the contribution of manual segmentation and the need for a combination of approaches.

### Differential effect of early hypoxia-ischemia on hippocampal subregions

Our observation that the CA-DG were the most affected hippocampal subregions in patients with developmental amnesia is congruent with documented reports of greater sensitivity of this subregion to hypoxia-ischemia. It has long been known that hypoxia and ischemia result in selective neuronal death in the CA1 field^21^, although the reason for this selectivity is still debated^23^. Neurohistological studies of patients who suffered ischemic episodes in adulthood have reported cases with neural cell loss restricted to the CA1 region bilaterally whilst the CA1’ region of the uncus, the subiculum, CA3 and surrounding cortices were all preserved^25,55^. High-resolution magnetic resonance imaging showed that CA1 was always affected in patients with hippocampal damage^24^.

In contrast, the uncus showed a low sensitivity to hypoxia in our patient cohort which could be a reflection of the distinct vascularization of this hippocampal subregion. The branches of the posterior cerebral artery (PCA) coming from the vertebral artery irrigate the posterior part of the hippocampus, while the branches of the anterior choroidal artery (AchA) originating from the internal carotid artery irrigate the uncal portion of the hippocampus^56–58^. High-resolution angiography and anatomical studies have highlighted that the anterior hippocampal region could receive a mixed blood supply (from both the PCA and the AchA) in about half of the individuals^56,59^. A mixed blood supply could provide more vascular reserve and lower vulnerability to neuronal injury and atrophy^60^. The uncus, which receives a distinct blood supply compared to the rest of the hippocampus, might thus exhibit greater resistance to the adverse effects of hypoxic-ischemic events than the other hippocampal regions.

### Uncus and recall memory

The residual uncus could be partly functional in patients with developmental amnesia and could be recruited for recall memory. We reported here that the volume of the uncus correlated negatively with recall scores as well as working memory performance in patients with developmental amnesia. In the WAIS/WISC test, Working Memory Index is made up of two subtests (Digit Span and Letter Number Sequencing) both requiring verbal recall within the span of working memory. The function of the uncus is not well understood mainly because its structure-function mapping remains to be determined in rodents^61^, but in humans, the uncus has been proposed to be involved in the long-term recall of scenes^62^. Connectivity studies in nonhuman primates have shown that the uncus is less connected to other regions of the hippocampus, and that it has its own associational system of connections^63^. Notably, the uncus is also the only hippocampal region reciprocally connected to its contralateral counterpart^43^. The uncus’ unique connectivity could allow this region to support some information processing despite severe damage to the other hippocampal areas.

### Working memory

We reported here, in addition to an absence of working memory impairment, a negative association between uncus volume and working memory performance in patients with developmental amnesia. The more established view proposes that the hippocampus plays an important role in episodic memory, spatial cognition, and navigation, while it is not critically contributing to working memory^64,65^. But an alternative view proposes that the hippocampus may contribute to any memory task, including working memory^66,67^, in particular in supporting high-precision binding^68^. This latter possibility is challenged however by the fact that patients with developmental amnesia presenting with severe hippocampal damage have repeatedly shown unimpaired working memory performance^69–71^. Recent work extended these findings by showing that high resolution working memory performance was unimpaired, and even often superior, relative to control group in a patient with developmental amnesia^72^. Our observation of a negative correlation between uncus volume and working memory performance suggests that the perirhinal, entorhinal, and parahippocampal cortices normally involved in the binding of information in the short term^73^ may be subject to plastic changes in patients with developmental amnesia and that these plastic changes may depend on the degree of hippocampal damage. The possibility that the hippocampal damage in patients with developmental amnesia may not influence working memory performance directly, but indirectly via reorganization of the MTL circuit is explored below.

### Functional hypotheses

Experimental studies in nonhuman primates have reported similar counterintuitive findings to those reported in the present study. Monkeys with small hippocampal lesions exhibit greater memory loss on a Delayed Nonmatching-to-Sample (DNMS) task than that found in animals with greater damage^74^. Furthermore, a positive correlation between the extent of damage to the hippocampus and scores on a DNMS task has been reported in experimentally lesioned monkeys^75^. Several hypotheses have been proposed to explain why discrete lesions result in greater memory impairment^74,76,77^. However, it is difficult to relate these findings obtained following adult experimental lesions in the monkey to observations in patients with early hippocampal damage whose circuits might have undergone massive compensation. One hypothesis to explain our observations is that the level of hippocampal damage would likely have a strong direct influence on the organization of the medial temporal lobe memory circuits in the adult. Following an early event of hypoxia-ischemia, greater hippocampal damage might induce greater compensatory reconfigurations in the neural circuits and enable other structures, in particular the surrounding cortical areas normally recruited by the hippocampus, to assume important aspects of memory function. In contrast, when the hippocampus is only partially damaged, the information flow would remain present in the hippocampus and could result in incomplete, and possibly disruptive information processing. Our observation could thus reveal the existence of multiple, redundant routes within the residual hippocampal subregions and surrounding cortical areas in patients with developmental amnesia^78^. These parallel circuits could compete for the control of behaviour and disrupt memory performance^77^.

Our findings are consistent with the hypothesis that a close relationship can be evidenced between the volume of remaining hippocampal subregions and the severity of cognitive deficits in patients with hippocampal damage, in the case of patients with developmental amnesia this relationship was inverse. Our observations also provide further support for the model of hierarchical organization of cognitive memory^79^ by proposing that surrounding cortical areas normally involved in the encoding of associative relations between decontextualized information^73^ could reorganize to contribute to hippocampal function following early hippocampal damage. The limits of this reorganization, however, may be determined by the extent of damage to the hippocampal subregions.

Evidence for such reorganization has already been reported in a nonhuman primate model of early hippocampal damage. Monkeys with early bilateral hippocampal lesion can learn new spatial relational information in striking contrast to monkeys with adult lesion^80,81^. Interestingly, in these monkeys with early hippocampal lesions, the surrounding cortical areas were largely preserved and were shown to contribute to spatial learning in the absence of functional hippocampal circuits^82^. Also, a recent report showed that the number of immature neurons in the entorhinal and perirhinal cortices was higher in monkeys with early hippocampal lesions than in controls^83^. This increase in the number of immature neurons was paralleled by an increase in the number of mature neurons in the entorhinal cortex of neonate-lesioned monkeys compared to controls. These structural changes may contribute to some functional recovery after early hippocampal injury^83^.

One could speculate that the functional reorganization of the perihippocampal structures (normally devoted to the processing of decontextualized information) to compensate hippocampal function could, in turn, benefit the processing of semantic memory and lead to better context-free memory performance in patients (see:^84^). Indications of increased memory performance have indeed been reported in two different patients with developmental amnesia. Jonin *et al*.^27^ studied patient ‘KA’ presenting with 55% hippocampal volume reduction and preserved perihippocampal structures. While patient KA displayed few, if any, residual episodic abilities, this patient was able to accurately retrieve semantic memories, and show evidence of superior or even very superior access to these memories compared to controls. Another patient with developmental amnesia, patient ‘Jon’, has often produced response accuracy levels that are at least higher than those of control participants in previous experimental explorations of visuospatial working memory^69,71,72^. Jon was at least numerically superior to the control mean on recognition-based measures of shape-color binding^71^. In measures of color-location memory, Jon’s recognition accuracy matched the highest achieving control participant, while his reconstruction performance was superior to 6 of the 7 controls^69^. Across four tasks measuring binding between color and orientation, or color and location using simultaneous or sequential presentation of stimuli, Jon’s response accuracy was high, and always numerically superior to the control mean^72^.

These observations from single patients studies together with the present findings, obtained in a large cohort of patients with developmental amnesia, support the hypothesis of a reorganization of the relatively preserved perihippocampal areas following early hippocampal damage.

### Conclusions

Our results have important implications for understanding the organization of the memory circuit following early hippocampal injury. These observations highlight the counterintuitive finding that in patients with developmental amnesia greater hippocampal damage can be associated with better performance in different cognitive processes, such as recall memory, verbal reasoning, working memory, and speed of information processing. In contrast, less severe damage, leaving behind relatively preserved hippocampal subregions, can lead to more pronounced cognitive deficits. These unexpected findings can disentangle the functional organization of the medial temporal lobe memory system, and attribute a central role in the compensation of cognitive memory to extra-hippocampal structures in patients with early hippocampal damage.

## Data availability

Anonymized participant data will be available upon reasonable request.

## Acknowledgements

We thank the late Mortimer Mishkin, Laboratory of Neuropsychology NIMH, Bethesda, MD, for his unfailing inspiration, encouragement and support enabling the realization of this research. We also thank the participants and their families for their support of our research.

