## Supplementary information for "Anatomo-functional changes in neural substrates of cognitive memory in developmental amnesia: Insights from automated and manual MRI examinations"

**The *Doors and People Test.*** The test consists of four subtests, two assessing recognition and two assessing recall, and, within each of these pairs, one assessing visual ability and the other, verbal ability. The recognition and recall subtests are matched for difficulty. The four subtests, which are labeled ‘People’, ‘Shapes’, ‘Names’, and ‘Doors’, measure verbal recall, visual recall, verbal recognition, and visual recognition, respectively. For verbal recall (‘People’ subtest), four photographs each depicting an individual together with their printed name and occupation were presented on separate cards. After viewing the fourth picture, participants were asked to recall each name cued by their profession. This procedure was repeated until all four names were correctly recalled, or for a maximum of three presentations. Similarly, for Visual recall (‘Shapes’ subtest), participants copied each of four simple line drawings. They then tried to draw the four shapes from memory. This procedure was repeated until all four shapes were correctly recalled, or for a maximum of three presentations. The Verbal recognition subtest (‘Names’ subtest) consisted of two study-test blocks. In the study phase of the first block, 12 female first names and surnames were presented on separate cards for 3 s each, and the experimenter read them aloud. Immediately thereafter, participants saw 12 lists of four names, each list presented on a separate card, and asked in each case to select the name from the study list. The same procedure was repeated in a second block, this time consisting of male names, but with the foils and the names on each test list differing from the study list in only one syllable of the surname. Finally, the Visual recognition subtest (‘Doors’ subtest) also consisted of two study-test blocks. In the study phase of the first block, participants viewed photographs of 12 doors, each presented on separate sheets accompanied by an appropriate label. Immediately thereafter, participants viewed 12 arrays of four doors, each on a separate sheet, and tried to identify the door from the study list. This same subtest was repeated with a second block consisting of 12 photographs of doors presented in exactly the same way as the first study-test block, but with foils that are more similar to the doors on the study list than is the case on the first block. Delayed (cued) verbal recall was tested after completion of the visual recognition test, and, similarly, delayed (cued) visual recall was tested after completion of the verbal recognition test. As all four subtests were designed to be equally difficult based on the performance of a large group of healthy participants, the scores of our patients on all four subtests could be directly compared.

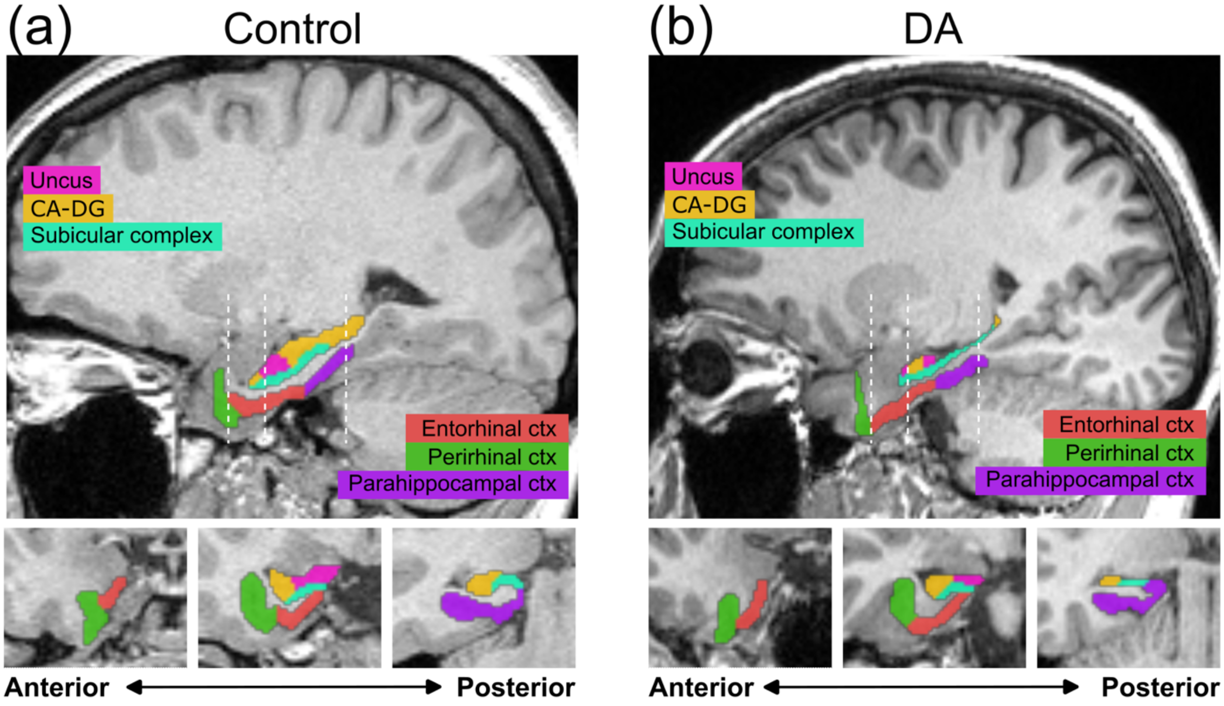


**Supplementary Figure 1 Manual segmentation of the hippocampal subregions and surrounding cortical areas.** (**a**) 1.5T-MRI scan of a 25-year-old female control participant. (**b**) 3T-MRI scan of a 25-year-old male patient with developmental amnesia presenting a 50% volume atrophy of the hippocampus. Sagittal and coronal views of the right hemispheres. CA-DG: orange; Uncus: magenta; subicular complex: cyan; entorhinal cortex: red; perirhinal cortex: green; parahippocampal cortex: purple.

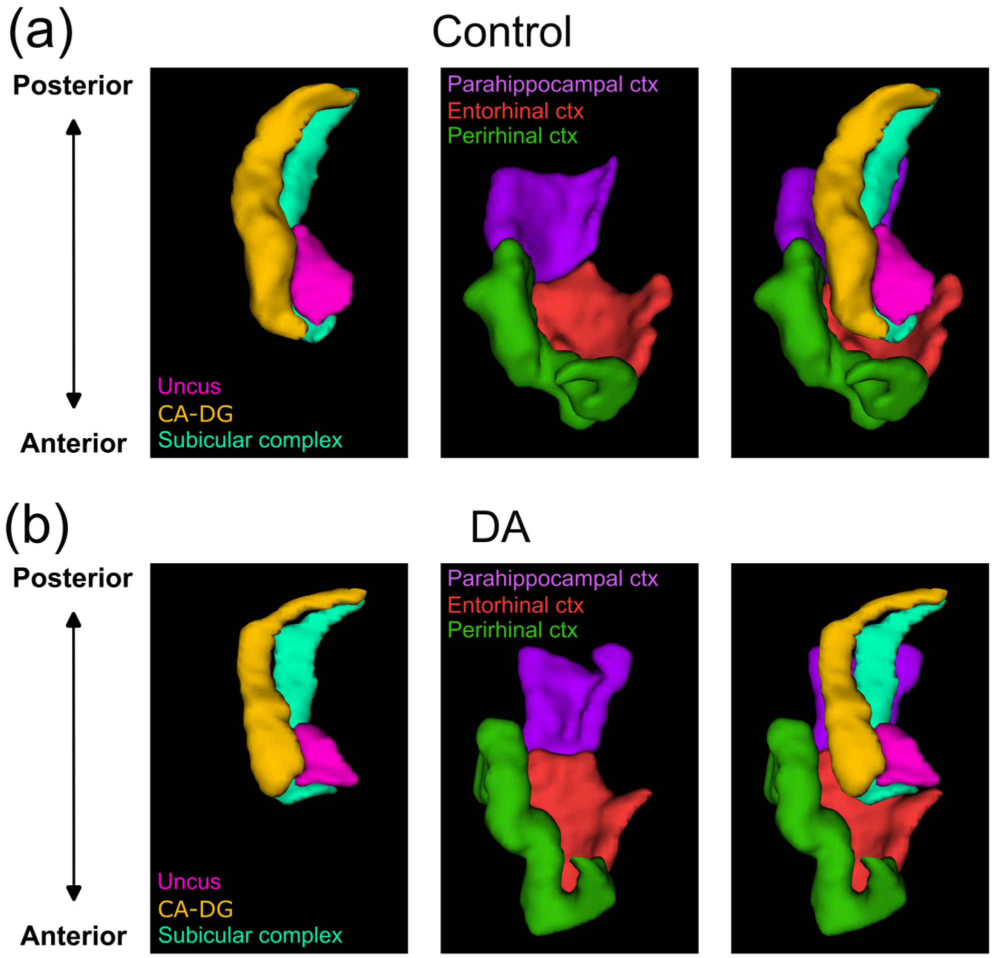


**Supplementary Figure 2 3D representations of the hippocampal subregions and cortical areas examined in this study, viewed from an antero-dorsal perspective.** (**a**) Control participant (25-year-old female). (**b**) Patient with developmental amnesia presenting a 50% volume atrophy of the hippocampus (25-year-old male). Views of the right hemispheres. CA-DG: orange; Uncus: magenta; subicular complex: cyan; entorhinal cortex: red; perirhinal cortex: green; parahippocampal cortex: purple.

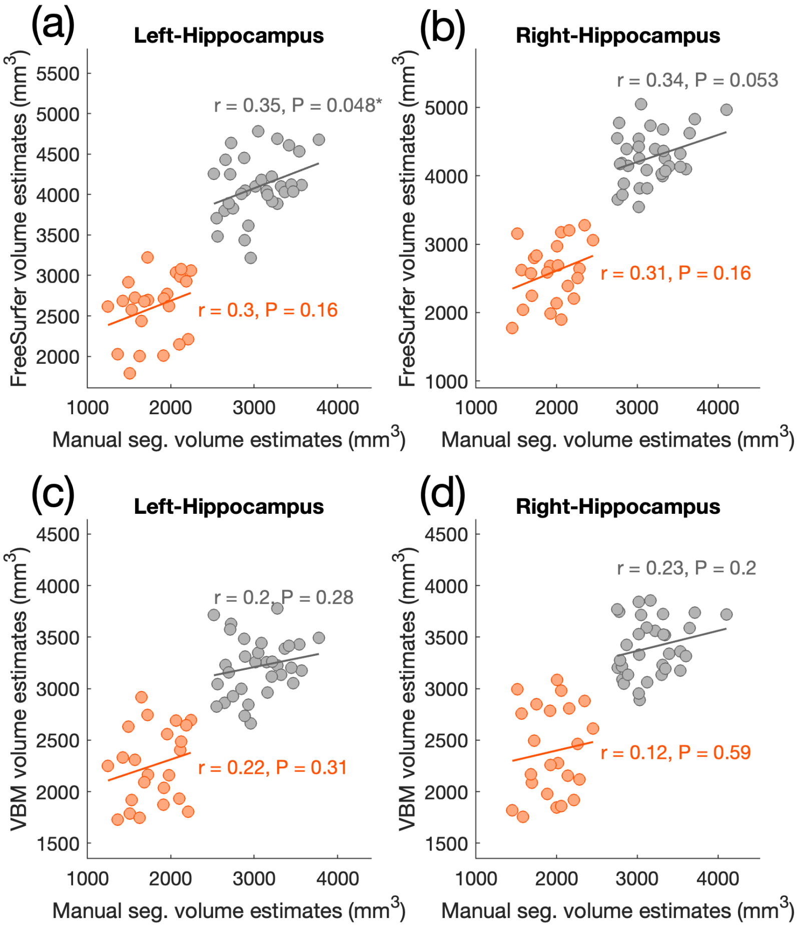


**Supplementary Figure 3 Correlations between the left and right hippocampal volume estimates obtained by VBM, FreeSurfer, and manual segmentation protocols.** Volume estimates by FreeSurfer (**a-b**) and voxel-based morphometry (**c-d**) plotted against manual segmentation estimates in control (grey; $N$ = 32) and developmental amnesia (orange; $N$ = 23) groups. Pearson's correlation coefficient. Manual segmentation volumes estimates are corrected for intracranial (ICV) volume.*: $P < .05$.

**Supplementary Table 1** Aetiology and neuroradiological assessment of MRI scans of patients with developmental amnesia (*N* = 23).

| | Age at onset | Aetiology | HPC | MTL | Fornix | Th | MB | CC | BG | PWM | Cb | LV | Other |
|---|---|---|---|---|---|---|---|---|---|---|---|---|---|
| DA01 | P | Central cyanosis, TGA, collapsed lung, ventilated | s | N | s | N | s | no | N | N | N | N | |
| DA02 | P | Prematurity (28w), intracranial haemorrhage, RDS | s | N | s | N | s | no | N | Focal abn bilateral | s | N | |
| DA03 | P | Prematurity (26w), intubated, ventilated, RDS | s | N | s | N | s | no | N | Mild PVL | s | N | |
| DA05 | 9y | Respiratory arrest at age 9y, ventilation, TLE from age 17y | s | N | N | N | N | no | N | N | N | N | |
| DA06 | 9y | Diabetes type I at age 8y, hypoglycaemic episode at age 9y, acute attack during sleep at 15y | s | N | s | N | N | yes | N | Focal abn bilateral | N | dilated | |
| DA08 | P | Severe asphyxia at birth, hypoxic-ischemic encephalopathy | s | N | s | N | s | no | N | N | s | N | |
| DA09 | P | Resuscitation & ventilation, RDS | s | N | very s | N | s | no | N | N | N | N | |
| DA10 | 12y | Tectal plate tumor, hydrocephalus, meningitis at age 12y, intraventricular bleeding, RDS | s | N | very s | N | s | yes | N | N | N | s (shunted) | [a] |
| DA11 | P | Prematurity (32w), intubated, ventilated, RDS | s | N | very s | N | s | yes | N | N | N | Mild dilated | |
| DA12 | P | Congenital heart defect, pneumonia, respiratory arrest at age 11w, TLE from age 9y | s | N | s | N | s | yes | N | N | N | N | [b] |
| DA13 | P | Cardiac arrest, shoulder dystocia, resuscitation, ventilation, RDS | s | N | very s | N | s | yes | N | PVL, Focal abn bilateral | s | dilated | |
| DA14 | P | Second-born twin with cord around his neck, RDS | s | N | s | N | s | yes | N | N | N | N | |
| DA15 | P | Congenital heart disease, cardiac failure at age 2y, heart transplant at age 3y | s | N | s | N | s | no | s on L | Focal abn L | N | Dilated; also L>R | [c] |
| DA16 | P | Fetal distress, cord around his neck, RDS | s | N | very s | N | s | no | N | N | N | N | |
| DA17 | P | RDS at birth, congenital hypothyroidism | s | N | very s | N | s | yes | N | N | N | N | [d] |
| DA18 | P | TGA, RDS | s | N | s | N | s | no | N | Focal abn R | N | N | |
| DA19 | P | Second born twin, resuscitation, ventilation, RDS | s | N | s | N | s | no | N | N | N | N | |
| DA20 | P | Prematurity (35w), intraventricular bleeding, resuscitation, RDS | s | N | s | N | s | no | N | N | N | N | |
| DA21 | P | Meconium aspiration syndrome, RDS | s | N | N | N | s | yes | N | N | N | N | |
| DA22 | P | Diaphragmatic hernia, lung atrophy, ECMO for 2 weeks | s | N | s | N | s | no | N | N | N | mild dilated R | |
| DA23 | P | Cardiac arrest, shoulder dystocia, RDS | s | N | s | N | s | no | N | N | N | N | |
| DA24 | P | Respiratory problems, ECMO for 3w, diabetes at age 6y | s | N | s | N | s | yes | N | N | N | N | [e] |
| DA25 | P | Prematurity, low birthweight, neonatal hypertension, perinatal pneumothorax | s | N | s | N | s | yes | N | N | N | N | |

[a] Right frontal ventricular catheter; Left frontal shunt track
[b] Minor cerebellar cortical injury
[c] Focal Left hemisphere cortical and white matter injury
[d] Focal abnormalities Left Claustrum
[e] Small anterior commissure
Abbreviations: abn, abnormalities; BG, Basal ganglia; Cb, Cerebellum; CC, splenium of the corpus callosum smaller in size than the genu, in contrast to the reverse pattern seen in the healthy population (yes/no); ECMO, Extracorporeal membrane oxygenation; HPC, hippocampus; L, left; LV, Lateral ventricle; MB, mammillary bodies; MTL, medial temporal lobe; N, normal; P, perinatal; PVL, periventricular leukomalacia; PWM, Periventricular white matter; R, right; RDS, Respiratory distress syndrome; s, small; TGA, transposition of the great arteries; Th, Thalamus; TLE, Temporal lobe epilepsy

**Supplementary Table 2** Volume of the hippocampus, hippocampal subregions and surrounding cortical areas in controls ($N$ = 32) and patients with developmental amnesia ($N$ = 23). ROI were manually segmented on MRI scans. Volumes are calculated as the average of left and right hemisphere volumes. ICV: intracranial volume.

| | | | | | ICV-corrected volumes ($mm^3$) | | | | | | |
|---|---|---|---|---|---|---|---|---|---|---|---|
| Code | Gender | Age (years) | MRI type | ICV ($mm^3$) | Hippocampus | Uncus | CA-DG | Subicular complex | Entorhinal | Perirhinal | Parahippocampal |
| C01 | M | 18.5 | 3T | 1'479'650 | 3'245 | 412 | 1'872 | 960 | 1'543 | 2'207 | 1'600 |
| C02 | M | 14.0 | 3T | 1'291'510 | 3'337 | 432 | 1'824 | 1'081 | 1'413 | 2'230 | 1'794 |
| C03 | F | 18.4 | 3T | 1'336'220 | 3'551 | 517 | 1'901 | 1'133 | 1'493 | 2'214 | 1'731 |
| C04 | M | 18.8 | 3T | 1'601'470 | 3'119 | 365 | 1'734 | 1'021 | 1'406 | 2'169 | 1'700 |
| C05 | M | 17.9 | 3T | 1'325'580 | 3'463 | 474 | 1'891 | 1'098 | 1'519 | 2'363 | 1'774 |
| C06 | F | 18.2 | 3T | 1'429'700 | 3'166 | 363 | 1'775 | 1'028 | 1'475 | 2'271 | 1'712 |
| C07 | M | 8.3 | 3T | 1'531'850 | 3'346 | 483 | 1'712 | 1'151 | 1'536 | 2'198 | 1'661 |
| C08 | F | 8.5 | 3T | 1'292'440 | 2'658 | 406 | 1'439 | 812 | 1'466 | 2'061 | 1'562 |
| C09 | M | 13.0 | 1.5T | 1'521'740 | 3'940 | 524 | 2'036 | 1'380 | 1'488 | 2'383 | 1'761 |
| C10 | M | 32.0 | 1.5T | 1'535'580 | 3'291 | 573 | 1'582 | 1'136 | 1'514 | 2'356 | 1'810 |
| C11 | F | 38.0 | 1.5T | 1'252'760 | 2'995 | 646 | 1'371 | 978 | 1'609 | 2'380 | 1'583 |
| C12 | F | 20.0 | 1.5T | 1'554'750 | 3'628 | 619 | 1'841 | 1'168 | 1'593 | 2'287 | 1'670 |
| C13 | F | 25.0 | 1.5T | 1'367'350 | 3'380 | 612 | 1'756 | 1'011 | 1'575 | 2'326 | 1'656 |
| C14 | F | 35.0 | 1.5T | 1'344'590 | 3'116 | 479 | 1'607 | 1'030 | 1'500 | 2'363 | 1'627 |
| C15 | M | 28.0 | 1.5T | 1'297'970 | 3'410 | 653 | 1'734 | 1'024 | 1'564 | 2'344 | 1'543 |
| C16 | M | 14.8 | 1.5T | 1'622'852 | 2'782 | 482 | 1'376 | 924 | 1'524 | 2'378 | 1'505 |
| C17 | F | 20.6 | 1.5T | 1'380'540 | 3'534 | 571 | 1'724 | 1'239 | 1'492 | 2'207 | 1'667 |
| C18 | M | 15.9 | 1.5T | 1'585'506 | 3'053 | 425 | 1'402 | 1'226 | 1'407 | 2'253 | 1'524 |
| C19 | M | 17.1 | 1.5T | 1'480'457 | 3'220 | 490 | 1'672 | 1'058 | 1'402 | 2'100 | 1'384 |
| C20 | F | 21.3 | 1.5T | 1'407'860 | 3'012 | 422 | 1'504 | 1'086 | 1'552 | 2'360 | 1'675 |
| C21 | F | 13.7 | 1.5T | 1'328'490 | 3'093 | 398 | 1'611 | 1'083 | 1'561 | 2'430 | 1'643 |
| C22 | F | 8.2 | 1.5T | 1'587'250 | 2'878 | 459 | 1'435 | 984 | 1'558 | 2'310 | 1'673 |
| C23 | F | 11.7 | 1.5T | 1'414'830 | 2'882 | 365 | 1'391 | 1'127 | 1'455 | 2'214 | 1'606 |
| C24 | F | 14.2 | 1.5T | 1'629'350 | 2'749 | 438 | 1'446 | 865 | 1'612 | 2'262 | 1'557 |
| C25 | F | 8.3 | 1.5T | 1'323'540 | 2'790 | 467 | 1'277 | 1'046 | 1'519 | 2'331 | 1'539 |
| C26 | M | 9.3 | 1.5T | 1'437'290 | 2'732 | 344 | 1'432 | 957 | 1'346 | 2'286 | 1'579 |
| C27 | F | 10.9 | 1.5T | 1'307'990 | 2'717 | 401 | 1'296 | 1'020 | 1'554 | 2'282 | 1'629 |
| C28 | M | 9.8 | 1.5T | 1'526'960 | 3'099 | 545 | 1'543 | 1'011 | 1'475 | 2'327 | 1'590 |
| C29 | M | 11.3 | 1.5T | 1'366'120 | 2'815 | 357 | 1'500 | 959 | 1'479 | 2'247 | 1'559 |
| C30 | F | 20.5 | 1.5T | 1'226'164 | 2'949 | 449 | 1'430 | 1'070 | 1'390 | 2'099 | 1'450 |
| C31 | M | 20.8 | 1.5T | 1'591'079 | 2'731 | 435 | 1'372 | 924 | 1'470 | 2'212 | 1'633 |
| C32 | M | 23.0 | 1.5T | 1'577'280 | 2'856 | 385 | 1'370 | 1'102 | 1'488 | 2'221 | 1'532 |
| DA01 | M | 16.0 | 1.5T | 1'325'550 | 1'928 | 428 | 930 | 570 | 1'414 | 2'497 | 1'646 |
| DA02 | F | 17.0 | 1.5T | 1'070'390 | 1'984 | 454 | 889 | 641 | 1'453 | 2'176 | 1'596 |
| DA03 | M | 40.0 | 3T | 1'272'340 | 1'824 | 281 | 902 | 641 | 1'384 | 2'174 | 1'660 |
| DA05 | F | 36.0 | 1.5T | 1'225'590 | 2'100 | 445 | 857 | 798 | 1'588 | 2'187 | 1'548 |
| DA06 | M | 32.0 | 3T | 1'466'940 | 1'873 | 405 | 770 | 698 | 1'416 | 2'042 | 1'451 |
| DA08 | M | 16.0 | 1.5T | 1'475'671 | 2'035 | 394 | 929 | 712 | 1'474 | 2'083 | 1'446 |
| DA09 | F | 17.0 | 1.5T | 1'023'350 | 2'210 | 432 | 1'045 | 734 | 1'494 | 2'287 | 1'676 |
| DA10 | F | 20.0 | 1.5T | 1'218'521 | 2'051 | 361 | 984 | 706 | 1'513 | 2'115 | 1'510 |
| DA11 | F | 22.0 | 1.5T | 1'315'940 | 1'804 | 298 | 927 | 578 | 1'485 | 2'297 | 1'709 |
| DA12 | M | 14.0 | 1.5T | 1'280'401 | 2'123 | 414 | 998 | 712 | 1'491 | 2'056 | 1'653 |
| DA13 | F | 28.0 | 1.5T | 1'232'000 | 1'611 | 394 | 704 | 513 | 1'422 | 2'222 | 1'706 |
| DA14 | M | 11.0 | 1.5T | 1'158'636 | 1'537 | 313 | 658 | 567 | 1'436 | 2'240 | 1'686 |
| DA15 | M | 9.0 | 1.5T | 1'424'720 | 2'223 | 489 | 1'084 | 649 | 1'458 | 2'379 | 1'480 |
| DA16 | M | 30.0 | 3T | 1'530'460 | 1'528 | 309 | 734 | 485 | 1'558 | 2'324 | 1'384 |
| DA17 | M | 25.0 | 3T | 1'511'710 | 1'573 | 241 | 789 | 543 | 1'548 | 2'120 | 1'775 |
| DA18 | F | 12.0 | 1.5T | 1'505'720 | 1'658 | 264 | 742 | 651 | 1'431 | 2'075 | 1'647 |
| DA19 | F | 27.0 | 1.5T | 1'231'010 | 1'473 | 317 | 683 | 473 | 1'441 | 2'401 | 1'580 |
| DA20 | F | 14.0 | 1.5T | 1'506'820 | 1'464 | 322 | 683 | 459 | 1'422 | 2'141 | 1'515 |
| DA21 | M | 8.0 | 3T | 1'528'660 | 2'061 | 407 | 1'074 | 580 | 1'608 | 2'313 | 1'614 |
| DA22 | M | 18.0 | 3T | 1'381'040 | 1'937 | 419 | 922 | 595 | 1'408 | 2'129 | 1'667 |
| DA23 | F | 13.7 | 3T | 1'373'440 | 2'136 | 318 | 1'126 | 692 | 1'484 | 2'283 | 1'542 |
| DA24 | M | 11.9 | 1.5T | 1'354'810 | 1'580 | 291 | 703 | 585 | 1'473 | 2'272 | 1'628 |
| DA25 | F | 10.5 | 1.5T | 1'366'950 | 2'289 | 321 | 1'146 | 821 | 1'353 | 2'292 | 1'498 |

**Supplementary Table 3** Doors and People subtests scores correlation with the volume of the hippocampal subregions in control ($n = 21$) and developmental amnesia ($N = 23$) groups. Pearson's correlation coefficient. *: $P < .05$ ; **: $P < .01$.

| | | Verbal recall (People) | Visual recall (Shapes) | Verbal recognition (Names) | Visual recognition (Doors) | RECALL | RECOGNITION |
|---|---|---|---|---|---|---|---|
| Control | Hippocampus | 0.12 | 0.15 | 0.24 | 0.2 | 0.18 | 0.26 |
| | Uncus | –0.12 | 0.21 | 0.018 | 0.36 | 0.06 | 0.22 |
| | CA-DG | 0.12 | 0.19 | 0.29 | 0.038 | 0.21 | 0.19 |
| | Subicular complex | 0.2 | –0.034 | 0.17 | 0.22 | 0.11 | 0.23 |
| Developmental Amnesia | Hippocampus | –0.014 | –0.0057 | 0.13 | –0.18 | –0.0094 | –0.057 |
| | Uncus | **–0.47 *** | **–0.55 **** | –0.26 | –0.36 | **–0.58 **** | –0.36 |
| | CA-DG | 0.13 | 0.16 | 0.35 | 0.035 | 0.17 | 0.19 |
| | Subicular complex | 0.097 | 0.12 | –0.0067 | –0.3 | 0.12 | –0.19 |

**Supplementary Table 4** Full-scale IQ index scores correlation with the volume of the hippocampal subregions in control ($n = 23$) and developmental amnesia ($N = 23$) groups. Pearson's correlation coefficient. *: $P < .05$.

| | | FULL-SCALE IQ | *Verbal comprehension* | *Perceptual reasoning* | *Working memory* | *Processing speed* |
|---|---|---|---|---|---|---|
| Control | Hippocampus | 0.24 | 0.02 | 0.36 | 0.27 | 0.25 |
| | Uncus | 0.16 | –0.059 | 0.31 | 0.12 | 0.15 |
| | CA-DG | 0.15 | –0.017 | 0.24 | 0.12 | 0.27 |
| | Subicular complex | 0.29 | 0.12 | 0.38 | **0.46 *** | 0.14 |
| Developmental Amnesia | Hippocampus | –0.21 | –0.14 | –0.2 | –0.16 | –0.37 |
| | Uncus | **–0.51 *** | **–0.42 *** | –0.38 | **–0.45 *** | **–0.5 *** |
| | CA-DG | 0.058 | 0.099 | –0.052 | 0.088 | –0.16 |
| | Subicular complex | –0.3 | –0.24 | –0.2 | –0.25 | –0.39 |
